# On the origins of phenotypic parallelism in benthic and limnetic stickleback

**DOI:** 10.1101/2022.10.02.510516

**Authors:** Laura L. Dean, Isabel Santos Magalhaes, Daniele D’Agostino, Paul Hohenlohe, Andrew D. C. MacColl

## Abstract

Rapid evolution of similar phenotypes in similar environments, giving rise to *in situ* parallel adaptation, is an important hallmark of ecological speciation. However, what appears to be *in situ* adaptation can also arise by dispersal of divergent lineages from elsewhere. We test whether two contrasting phenotypes repeatedly evolved in parallel, or have a single origin, in an archetypal example of ecological adaptive radiation: benthic-limnetic three-spined stickleback (*Gasterosteus aculeatus*) across species-pair and solitary lakes in British Columbia. We identify two genomic clusters across freshwater populations, which differ in benthic-limnetic divergent phenotypic traits and separate benthic from limnetic individuals in species pair lakes. Phylogenetic reconstruction and niche evolution modelling both suggest a single evolutionary origin for each of these clusters. We detected strong phylogenetic signal in benthic-limnetic divergent traits, suggesting they are ancestrally retained. Accounting for ancestral state retention, we identify local adaptation of body armour due to the presence of an intraguild predator, the sculpin (*Cottus asper*) and environmental effects of lake depth and pH on body size. Taken together, our results imply a predominant role for retention of ancestral characteristics in driving trait distribution, with further selection imposed on some traits by environmental factors.

## Introduction

Parallel occurrence of adaptive phenotypes across similar but geographically separate environments has long fascinated evolutionary biologists. There are two main mechanisms which can explain such a pattern. First, novel adaptive phenotypes may evolve rapidly and repeatedly in response to new ecological opportunity i.e. parallel ecological speciation [1]. Alternatively, an adaptive phenotype may arise in a single location and disperse into and or persist only in suitable environments [2-4]. Although these two mechanisms result in the same pattern, they reflect extremely different evolutionary histories: multiple evolutionary origins of the same phenotype vs a single origin. It is therefore necessary to determine which evolutionary history is responsible for apparent parallelism if we are to understand it. There are many definitions for parallel and convergent evolution [5, 6], but here we focus on whether similar phenotypic adaptations share an ancestral genetic basis.

Parallel ecological speciation may involve multiple *de novo* mutations, each of which may lead to a similar phenotype but by a slightly different mechanism. In this instance it is easy to conclude multiple independent origins. However, evolution is not linear but often reticulated, and, in many cases, parallel ecological adaptation may involve repeated reuse of long-standing genetic variation i.e. the same, potentially ancient mutation can be introduced to multiple independent populations via admixture [7]. In this case, parallel populations may be young, and have multiple origins, but the mutations responsible for adaptation are shared and may be much older. This scenario is extremely difficult to differentiate from a scenario in which parallel populations themselves have a single origin [8], but it is critical that we attempt to do so in order to understand the underlying processes that shape evolution.

The benthic-limnetic axis of stickleback in British Columbia (‘BC’), Canada, is an archetypal example of ecological divergence and speciation [9-14]. It separates bottom-dwelling, benthic individuals, which feed predominantly on macroinvertebrates, from pelagic fish, feeding mostly on zooplankton [15-17]. These two freshwater ecotypes are characterised by heritable differences in body size, shape, trophic morphology and body armour, which confer fitness advantages in their corresponding habitats [18-20]. In BC, stickleback occur both as sympatric benthic-limnetic species pairs and solitary populations that possess phenotypes along the benthic-limnetic axis [9, 21-24]. Previous work has identified patterns of parallelism in adaptive genomic divergence across benthic-limnetic species pairs, but closer genetic affinity within lakes at neutral markers [21, 22, 25]. This work has tentatively led to the conclusion that benthic and limnetic phenotypes evolved repeatedly and independently in multiple lakes [21, 25, 26]. However, gene flow occurs to some extent in all benthic-limnetic species pairs [27, 28], and even low levels of gene flow quickly erode genetic differences at neutral loci, making it impossible to separate patterns of recent *in situ* ecological speciation from those derived from secondary contact of much older independent lineages [29]. Little investigation has so far been conducted beyond the species pairs, which coexist in only a handful of lakes [30], but see [31]. Populations in solitary lakes have far less opportunity for gene flow and thus will likely give a more reliable estimate of the evolutionary history of benthic and limnetic ecotypes.

We investigate whether benthic-limnetic divergence in BC stickleback likely has a single or multiple evolutionary origins. We first characterise genomic and phenotypic divergence across populations and show that all freshwater individuals fall within one of two genomic clusters, one of which exhibits a more benthic phenotype, and the other, a more limnetic phenotype. We construct a maximum likelihood phylogeny using a stringently filtered dataset, removing all known QTL in stickleback, and test for phylogenetic signal in ecologically relevant phenotypic traits. We construct a microevolutionary adaptive landscape for the BC radiation using recently available niche modelling techniques [32] to identify the best fitting model of benthic-limnetic trait evolution. Finally, accounting for any phylogenetic signal, we test for relationships between phenotype and environment to detect signals of true ecological adaptation.

## Results

We collected stickleback, and environmental parameters from 21 lakes surrounding the Strait of Georgia, BC (Figure 1), including two species pair lakes, two coastal locations (representing putative marine ancestors) and 17 solitary freshwater lakes (Table S1). We collected phenotypic data for key benthic-limnetic divergent traits (methods) for approximately 32 individuals (mean = 31.5, SE = 2.4) and generated RAD-seq genomic data [13], for approximately 16 individuals (mean = 15.9, SE = 0.9), from each lake, 333 individuals in total. This resulted in a master genomic dataset of 12,756 SNPs, which was subject to further filtering for some analyses (Table 1).

**Table 1.** SNP datasets. Details of the SNP datasets used in genomic analyses.

| Dataset | N lakes | N individuals | N SNPs | LD thinned | Known QTL removed |
| --- | --- | --- | --- | --- | --- |
| dataset 1 | 21 | 333 | 12,756 | ✗ | ✗ |
| dataset 2 | 21 | 333 | 9668 | ✓ | ✗ |
| dataset 3 | 21 | 333 | 6215 | ✓ | ✓ |
| dataset 4 | 5 | 53 | 9668 | ✓ | ✗ |

N: number, LD thinned: SNPs with $r^2 > 0.2$ removed, Known QTL: SNPs identified as falling within known QTL regions [59].

**Figure 1.**
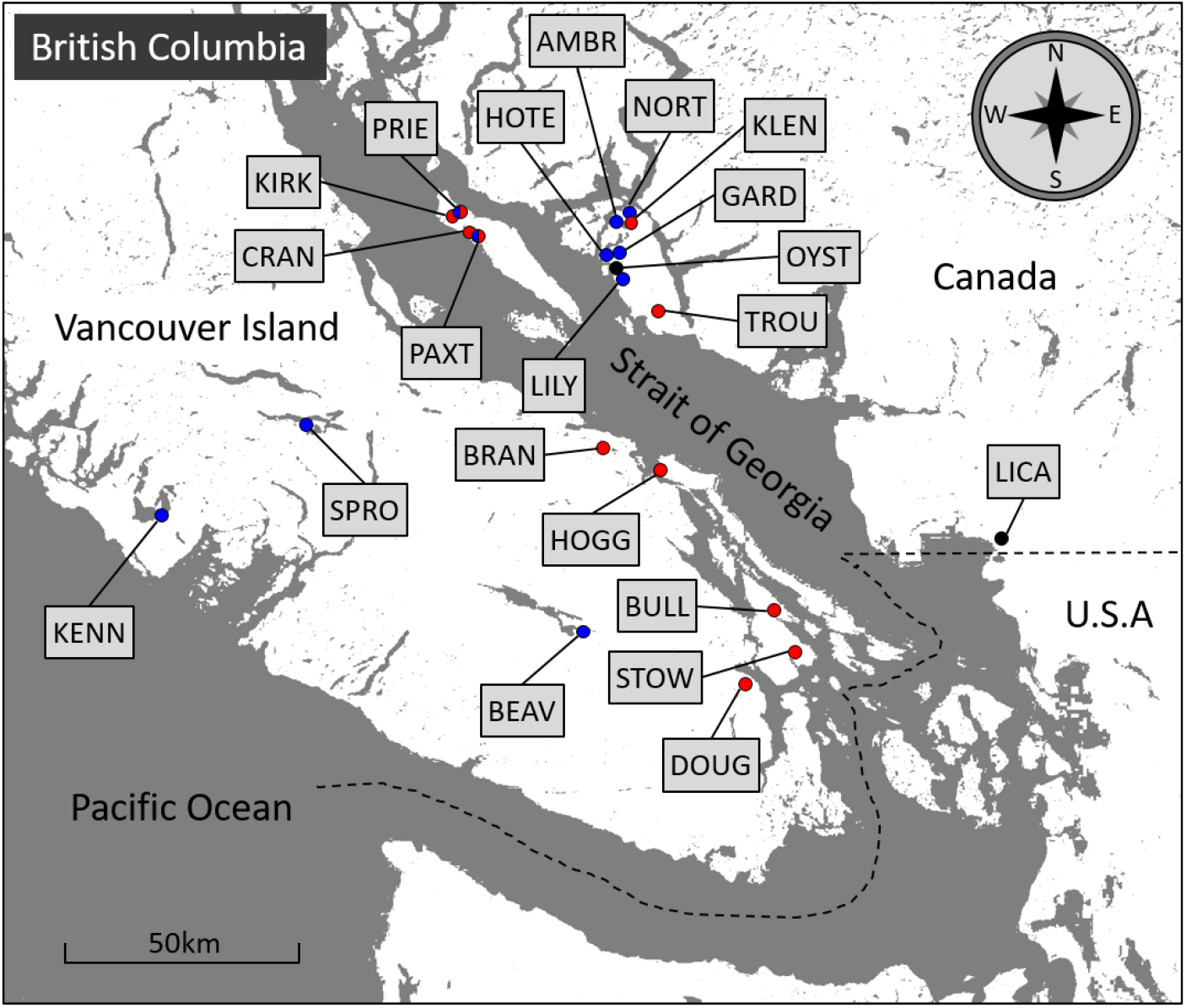
Map of sampling locations in British Columbia. Sample sites are indicated by circles and their associated lake ID. Black circles indicate marine populations, blue circles indicate populations in cluster 1 of our genomic analyses and red circles, cluster 2. Red and blue semi-circles indicate species-pair populations containing individuals from both clusters 1 and 2. The dashed line represents the border between Canada and the USA.

### Genomic divergence

We used two methods to quantify clustering within the genomic data. Firstly, a co-ancestry matrix in fineRADstructure [33] (dataset 1, 12,756 SNPs) identified two genomic clusters across all populations (Figure 2A), one incorporating the marine populations and approximately half of the freshwater populations (cluster 1), and the other comprising the rest of the freshwater lakes (cluster 2). Although marine populations formed part of cluster 1, they are considered separately here and in all further analyses because their presence in shallow coastal areas is transient and they likely represent the ancestral phenotypic state of all freshwater populations [7]. Secondly, we conducted a principal coordinate analysis (PCoA, dataset 2, 9668 SNPs). The same two broad genomic clusters (1 and 2) were identified by PCoA analysis, separating along PCo1 (9% of total variation, Figure 2B).

**Figure 2.**
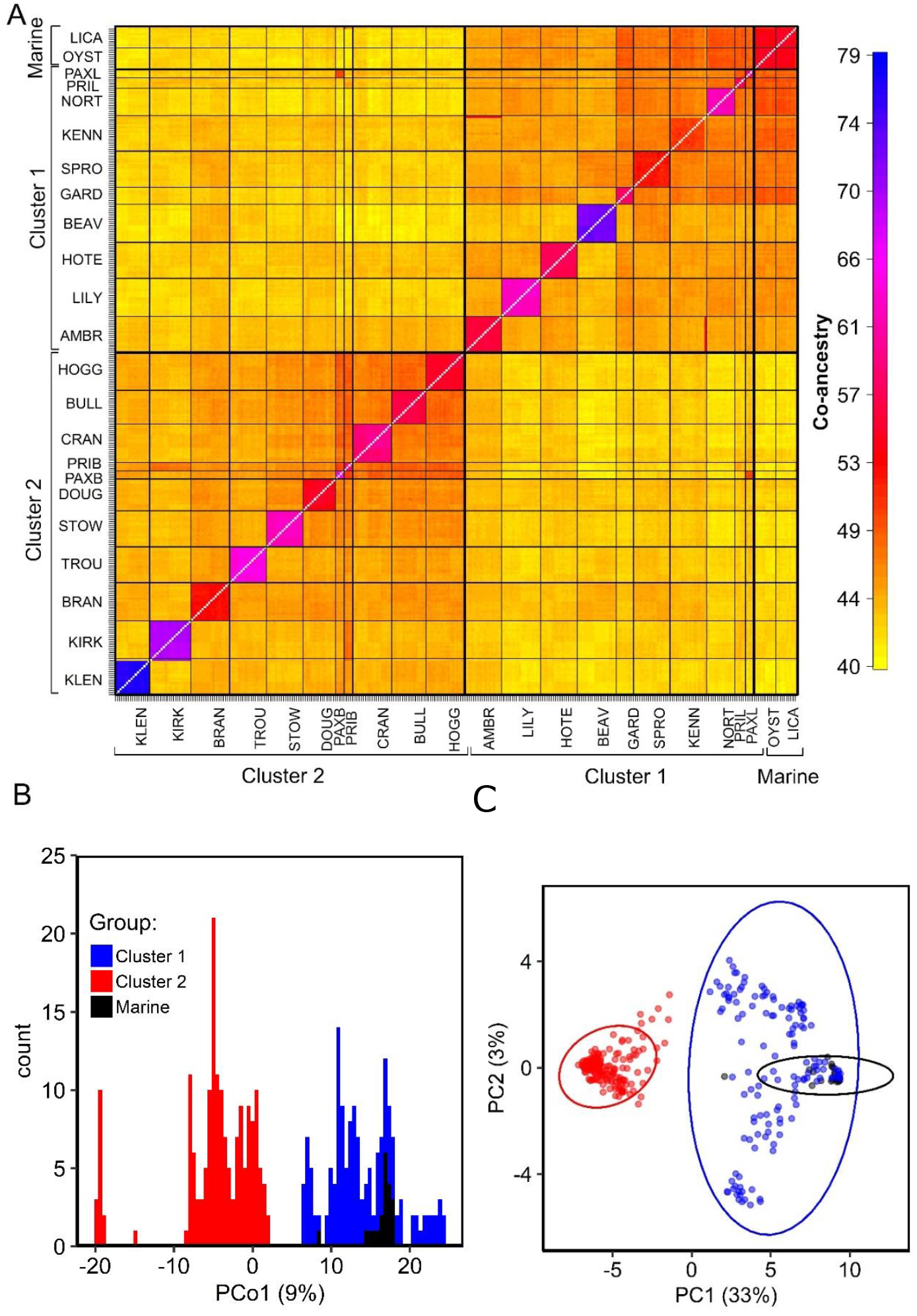
Genetic structure across BC stickleback populations. (A) Co-ancestry matrix of BC stickleback populations, using fineRADstructure. Thin black lines separate populations and thick black lines separate the broader genetic clusters. (B) Distribution of BC stickleback along the first principal coordinate of a genomic PCoA (dataset 2, 9668 SNPs). (C) Distribution of BC individuals in a PCA of 60 linked SNPs comprising linkage cluster 10, identified by LDna.

To further investigate the genomic properties of cluster 1 and 2 differentiation, we conducted a linkage disequilibrium network analysis (LDna) using the LDna R package (dataset 1, 12,756 SNPs). The resulting LD network contained twelve linkage clusters, each of which is likely a signature of a different evolutionary phenomenon [34]. Principal component analysis (PCA) on the SNPs from each cluster revealed a group of 60 SNPs, spread across 17 of the 21 chromosomes in the stickleback genome, associated with cluster 1 – cluster 2 separation. Of these 60 SNPs, 28 fell directly within genes (Table S2). Most other LD clusters only separated single populations from all others, likely reflecting local patterns of selection and drift and none of the LD clusters separated marine and freshwater adapted populations (Figures S1 and S2).

### Phenotypic divergence

To determine whether the genomic clusters differed phenotypically, we analysed differences in group means for important benthic-limnetic divergent phenotypic traits: weight, gill raker length and number, armour PC1 (increasing size of all armour variables and increasing lateral plate number, explaining 70.4% of body armour variation [methods]) and shape PC1 (describing shape changes associated with benthic and limnetic habitats, such as eye size, body depth and mouth length, explaining 23.2% of body shape variation [35, 36]). There were differences in phenotype between the three groups for all phenotypic traits (Table S3). For most traits, clusters 1 and 2 differed from marine fish, and for all traits except for body weight, clusters 1 and 2 differed from each other (Table S4, Figure 3A-E). Cluster 1 had a typically limnetic phenotype [35, 36] with a smaller size, longer, more numerous gill rakers, more body armour, a larger pelvis relative to spine length, and a more streamlined, slender body shape than cluster 2, which had a much more benthic phenotype (Figure 3A-E).

**Figure 3.**
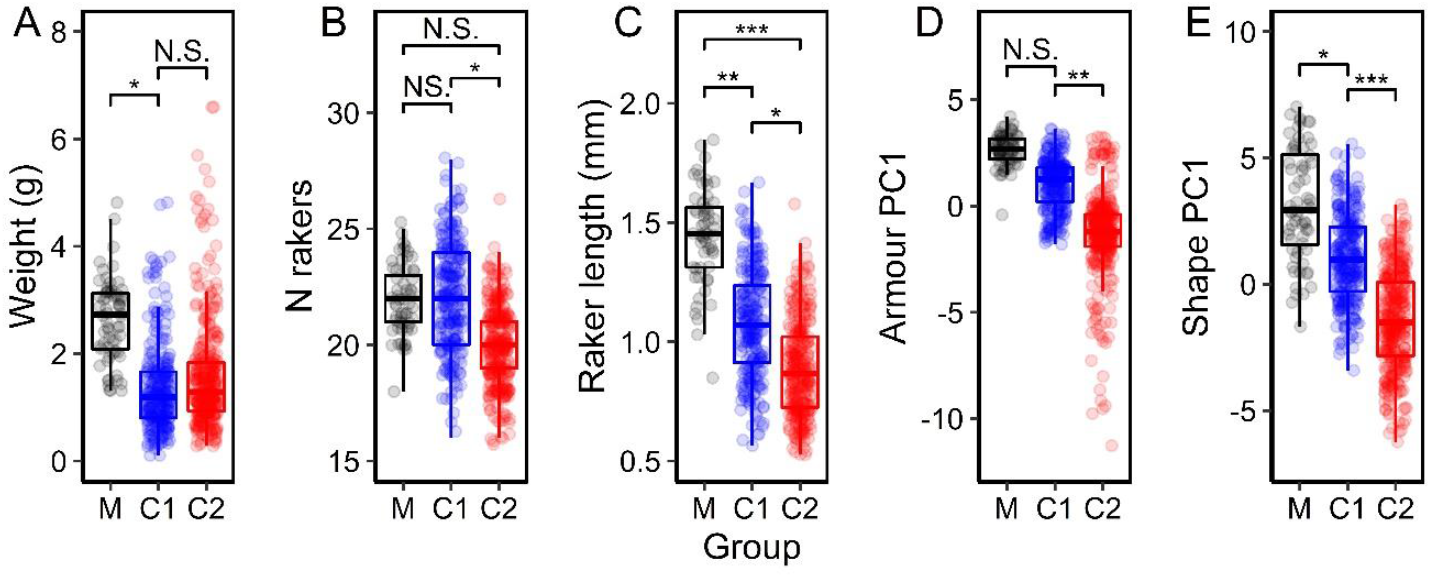
(A) – (E) Phenotypic differences between marine fish and two freshwater genetic clusters. Circles represent individuals. Abbreviations: M – marine, C1 – cluster 1, C2 – cluster 2. Brackets and asterisks indicate significance thresholds of post-hoc estimated marginal means tests between groups: NS. indicating *p* > 0.05, ** indicating *p* < 0.01 and *** indicating *p* < 0.001. All *p* values were adjusted for multiple comparisons.

### Phylogeny

Phylogenetic reconstruction for population-level genomic data can be notoriously problematic as numerous factors, including ongoing and / or historic gene flow can mask true phylogenetic signal in the data [37, 38]. To minimise bias in our analysis, we first filtered our master dataset (dataset 1, 12,756 SNPs) to remove all known QTL in stickleback (leaving 8351 SNPs, see methods), and then filtered for linkage disequilibrium (R^2^ > 0.2, leaving 6215 SNPs, dataset 3). We constructed a maximum likelihood phylogeny for all populations using RAxML. Consistent with the co-ancestry and PCoA analyses, the topology showed clusters 1 (more limnetic phenotype) and 2 (more benthic phenotype) at opposite ends of the tree, with marine fish most closely related to cluster 1 (Figure 4A). The two species-pair lakes both contained limnetic individuals whose closest relatives were in cluster 1 (PAXL and PRIL), and benthic individuals whose closest relatives were in cluster 2 (PAXB and PRIB, Figure 4A).

**Figure 4.**
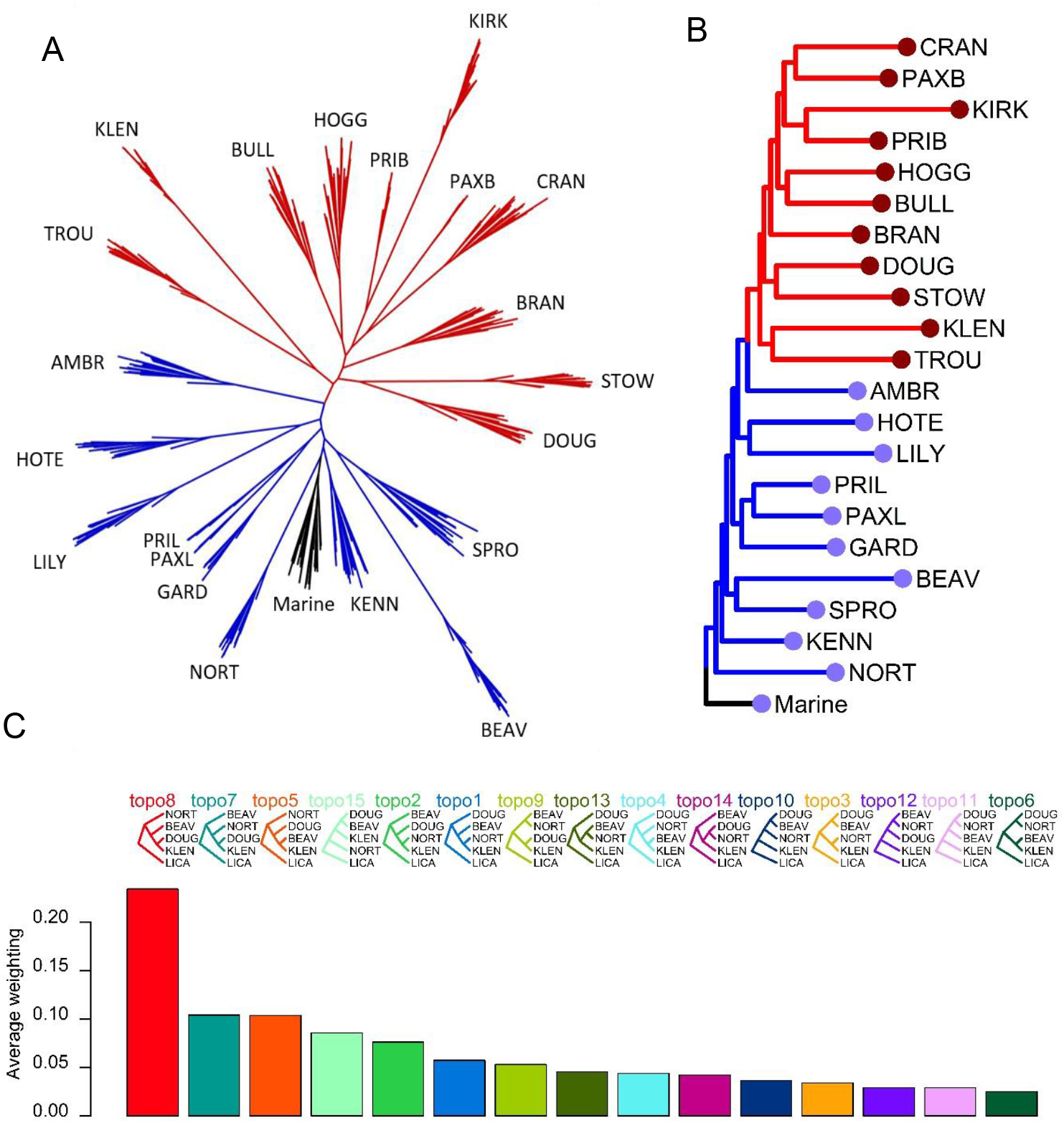
(A) Maximum likelihood phylogeny of 333 BC stickleback. Black indicates marine; blue, cluster 1 and red, cluster 2. (B) The same phylogeny as (A) with monophyletic populations collapsed into single tips. Branch colours in (B) denote the same as in (A). Coloured circles at branch tips represent two independent selection regimes, detected in the optimal model of niche evolution (R package: SURFACE). In both phylogenies species pairs are divided into benthic (PAXB, PRIB) and limnetic (PAXL, PRIL) populations. (C) Mean weightings for all possible topologies for four freshwater populations: two from cluster 1, NORT and BEAV; and two from cluster 2, DOUG and KLEN, with a single individual from the marine population LICA as the outgroup.

We also performed a topology weighting analysis on a subset of populations selected specifically to test the likelihood that the phenotypes associated with clusters 1 and 2 could have evolved repeatedly *in situ*. Topology weighting is a means by which to quantify relationships between taxa that are not necessarily monophyletic. It determines how support for each possible topology varies across the genome and allows quantification of the overall proportion of the genome which supports each possible tree. This allowed us to identify multiple highly supported phylogenies so that we could determine whether any of those with high support involved a model in which the two clusters arose more than once. It also allowed us to quantify what proportion of the genome supports our most likely topology, and how big the difference is between this and the level of support for the next most likely tree. To do this, we selected two populations from each of two locations approximately 100km apart and separated by the Georgia Strait. Each location contained a cluster 1 and 2 population occurring reasonably close together which, under a model based purely on geography, would be predicted to be more closely related. One marine individual was used as an outgroup. The topology with the highest weighting across all 50bp sliding windows (topology 8) was concordant with the maximum likelihood phylogeny, with the two cluster 2 populations (DOUG and KLEN) forming a monophyletic clade and each cluster 1 population splitting off earlier, deeper to the root (Figure 4C). The topology with the second highest weightings (topology 7) was also concordant, and just involved a switching of the order in which the cluster 1 populations split from the root. The simple geographical hypothesis, with the two pairs of populations nearest to one another being most closely related (topology 3) received very little support. The highest ranking topology had more than twice the proportional support that the second most likely topology had, suggesting that there is a strong genome-wide signal in favour of the maximum likelihood topology.

### Phylogenetic signal

If benthic and limnetic phenotypes had resulted from repeated, rapid adaptive divergence, phylogenetic signal (the tendency for more closely related individuals to share phenotypes) would be obscured, and trait distributions would instead mimic the adaptive landscape — i.e. variation in the relevant environmental characteristics. Therefore, we tested a null model that traits would be distributed randomly with respect to phylogeny, and an association of trait distribution with population-level relatedness was taken as evidence that benthic and limnetic niches were conserved from the ancestral state [39-41].

We estimated phylogenetic signal at the population level, using mean phenotypic trait data, and collapsing nodes in the phylogeny by population (with the two marine populations grouped into a single node, as they lacked monophyly), using the R package: PhyloSignal. We also tested five simulated traits that had no true association with phylogeny. We identified phylogenetic signal in all five real phenotypic traits: weight, gill raker number, gill raker length, armour PC1 and shape PC1 (*p-*values < 0.05, Table 2). None of the five randomly simulated traits showed phylogenetic signal (*p-*values > 0.05, Table 2).

**Table 2.** Phylogenetic signal in real and simulated phenotypic traits. Table shows estimates of phylogenetic signal (Pagel’s λ) and their associated *p*-values. *P*-values < 0.05 are highlighted in bold.

| Trait | Pagel's $\lambda$ | $p$-value |
| --- | --- | --- |
| Weight | 1.5881 | **0.0421** |
| Number of gill rakers | 2.1535 | **0.0010** |
| Mean gill raker length | 1.6492 | **0.0032** |
| Armour PC1 | 0.9667 | **0.0269** |
| Shape PC1 | 1.3647 | **0.0010** |
| Random 1 | 0.0001 | 1.0000 |
| Random 2 | 0.0001 | 1.0000 |
| Random 3 | 0.0001 | 1.0000 |
| Random 4 | 0.0001 | 1.0000 |
| Random 5 | 0.0001 | 1.0000 |

### Niche evolution modelling

Niche evolution modelling combines phylogenetic information with the distribution of phenotypic traits across the tree to identify the most likely number and location of selection regimes imposed across the whole phylogeny. It also identifies the number of instances of convergence (where the same regime appears multiple times across the tree). If the benthic phenotype had evolved repeatedly and independently across the phylogeny, niche evolution modelling should identify multiple instances of convergence of a benthic selection regime. We performed niche evolution modelling using the R package SURFACE [32]. We ran SURFACE using the same collapsed phylogeny and associated trait data that were used to estimate phylogenetic signal. The best fitting model involved two different selection regimes across the phylogeny (Figure 4B). The first included all marine and cluster 1 populations, and the second, all cluster 2 populations. The best fitting model included no instances of convergence between selection regimes, i.e. each independent regime appeared only once across the phylogeny.

### Relationship between phenotype and environment

To test for phenotype – environment relationships, we used linear mixed models, following a phylogenetic generalised least squares (PGLS) approach so that phylogenetic signal could be accounted for and fitted to the population means of phenotypic traits in R. Marine fish were excluded from all phenotype – environment modelling, because of the difficulty of measuring the environment of migratory marine fish. We found that freshwater fish had more body armour in the presence of sculpin (adjusted *p*-value < 0.05, Table S5), and fish were heavier in deeper lakes and lakes with a higher pH (adjusted *p*-values < 0.01 and < 0.05, respectively). Lake surface area and calcium concentration did not affect any aspect of phenotype and none of the environmental variables we measured affected the number of gill rakers, the length of gill rakers or shape PC1 (Table S5).

## Discussion

The repeated occurrence of similar phenotypes in geographically isolated but similar environments has several possible evolutionary explanations. Perhaps this pattern results from parallel ecological speciation, or maybe similar phenotypes have a single origin and have subsequently become widely dispersed into suitable habitats. It is impossible to separate these different models using only phenotypic data or small numbers of genetic markers, and remains difficult even with genomic data. Nevertheless, it is important that we attempt to do so, because of the consequences for our understanding of evolution. Parallel evolution has been implicated in the global adaptation of stickleback to freshwater [21], but recent analyses suggest that this is unlikely outside of the Eastern Pacific [42]. Furthermore, there are a number of other cases in which conclusions of parallel ecological speciation have been called into question by the confounding possibility of a single evolutionary origin followed by migration and gene flow [29].

We investigated the evolutionary origins of divergent phenotypes in a classic model system for adaptive radiation and ecological speciation. We find strong evidence for a monophyletic clade of stickleback with a benthic phenotype distributed across freshwater lakes in the southern Georgia Strait region of BC. The evidence strongly suggests that this clade has a single evolutionary origin, and is derived from a local ancestor with a limnetic phenotype.

The benthic clade also encompasses benthic fish from two benthic-limnetic species pairs. Our results are consistent with a single origin for benthic fish in BC rather than repeated independent evolution of the benthic phenotype in multiple lakes. This contradicts the currently favoured model for the evolution of benthic and limnetic stickleback in BC [21, 22, 25], highlighting the challenging nature of phylogenetic reconstruction at the population level.

Many factors, such as incomplete lineage sorting, hybridization, gene duplication, natural selection and recombination can lead to genealogical discordance in estimations of phylogenetic relationships [43]. Resolving the true relationships between divergent groups can therefore be challenging and require a large number of genetic markers. Much of the current research on benthic and limnetic stickleback in BC has been based only on mitochondrial haplotypes [25] or relatively small SNP sets [21], and has largely been restricted to species pairs. In species pairs, multiple QTL regions are repeatedly responsible for benthic adaptation [21, 44, 45], which is consistent with a single benthic origin, but neutral SNPs imply closer genetic affinity of benthics and limnetics within lakes [21], consistent with multiple independent origins. However, elevated levels of genetic similarity at neutral markers in species pairs would be expected even with low levels of gene flow and thus may not reflect true phylogenetic relationships [27]. Clearly, it is important also to consider relationships in solitary populations, where opportunity for gene flow is greatly reduced. Harer et al. [31] looked at both species pairs and solitary populations and identified considerable genomic parallelism in the former but not the latter. However, they define the benthic-limnetic spectrum in solitary populations solely using lake surface area, which correlates only weakly with phenotype, and does not account for differences in lake depth and the presence of specific predators, which also have a major influence on the availability of benthic and limnetic niches to stickleback [46].

We identify a monophyletic lineage within BC stickleback, which has a consistently benthic phenotype when compared with other freshwater or marine populations in BC. Some argument remains among evolutionary biologists about whether pervasive, genome-wide selection, can overwhelm the signal from other markers and obscure tree topologies [43, 47]. Sculpin (*Cottus asper*) are an intraguild predator of stickleback, which, when present, select for a more limnetic phenotype [48, 49]. Our results were consistent with this as we identified an effect of sculpin presence on body armour after accounting for phylogenetic signal in our analysis. However, we find the possibility that selection from sculpin obscures the true relationships between populations in our phylogeny unlikely for a number of reasons. Firstly, such a phenomenon is certainly possible in studies using only a small number of markers [50], but with many thousands of unlinked genetic markers, such as in this case, the probability of selection overwhelming the signal from neutral markers is very low [47, 51]. Secondly, recent modelling suggests that even with strong selection affecting 10-20% of markers, in most instances, phylogenetic inference remains robust to the effects of selection [52]. Selection from sculpin likely affects less than 2% of the genome [48] and thus is at least an order of magnitude smaller in effect size than would be necessary to obscure the true tree topology in this case. Furthermore, sculpin were present in some lakes containing the benthic clade and not in all lakes containing the limnetic clade. Thus, the genomic groups identified here do not simply mirror the occurrence of this selective agent, but rather represent a deeper set of ancestral relationships.

If the benthic phenotype in BC has a single origin, a clear question that requires explanation is how this ecotype spread across BC? Although the lakes containing stickleback in BC are not particularly widespread, they are physically separated by land or ocean, which likely makes dispersal a challenge for freshwater stickleback. We speculate that evidence for a large flood (∼500km^3^ of water) in the Fraser River valley, dated approximately to the end of the Pleistocene and caused by the failure of a large ice dam [53], could provide an explanation.

The estimated extent of the flood across the southern Georgia Strait is very similar to the current known distribution of benthic stickleback in BC, raising the tantalising possibility that it may have been responsible for the spread of the benthic lineage of stickleback from a palaeolake in the Fraser Valley, consistent with previous inference about the evolution of Eastern Pacific freshwater stickleback [42].

Our investigations have shown that the well-studied benthic-limnetic species pairs should be understood as part of a broader radiation along the benthic-limnetic axis in BC. We highlight the need to consider carefully all possible explanations for the occurrence of parallel phenotypes if we are to achieve a proper understanding of the evolutionary processes that mediate divergence. Stickleback are clearly capable of remarkably rapid ecological adaptation [54-58], but we have shown that the retention of ancestral characteristics can also be important in explaining the distribution of divergent phenotypes. This has significant implications for how we think about the process of evolution and raises the possibility that other model examples of *in situ* ecological adaptation may also result from dispersal rather than convergence.

## Acknowledgements

The authors would like to thank the Schluter lab for assistance with fieldwork and collection permits and Dolph Schluter for useful comment on previous drafts of the manuscript, Steve Vamosi for access to lake habitat data, Ann Lowe for assisting with the DNA extractions, and Muayad Mahmud and Becca Young for assistance with phenotyping. We would also like to thank Mark Ravinet for providing useful suggestions to improve the analyses and towards drafting the manuscript. Analyses of genomic data were conducted using the University of Nottingham’s on premise, high performance computer service, Augusta.

This work was funded by NERC grants NE/J02239X/1 and NE/R00935X/1.

## Methods

### Samples sites and environmental measurements

A total of 21 lakes surrounding the Strait of Georgia, BC, which were likely to vary substantially in the ecological niches they presented to stickleback (because of variation in environmental factors), were selected for sampling (see Table S1 for detailed sample site information and Figure 1 for a map of sampling locations). This included two lakes, Paxton (PAXT) and Priest (PRIE), known to contain benthic-limnetic stickleback species pairs [16, 26], and two coastal locations accessible from the sea, Oyster lagoon (OYST) and Little Campbell River (LICA), where marine fish are present during the spring breeding season.

The size and depth of a lake largely determine whether both benthic and limnetic habitats are present (in larger deeper lakes) or just benthic (in small, shallow lakes). Therefore, we measured the surface area (km^2^) using GoogleEarth and collected data on the mean depth (m) of each lake from either HabitatWizard [60] or from data collected in Vamosi [61], with permission. The presence of other fish species can also determine whether both, one or none of those niches are available to stickleback [61]. Many other fish species occur in BC, some of which are predators and/or competitors of stickleback. Cutthroat trout (*Oncorhynchus clarkii*) and rainbow trout (*Oncorhynchus mykiss*) are major intraguild predators of stickleback, but both occur in both the littoral and pelagic zones [62, 63] and do not eliminate either niche for stickleback and so are not considered further here. Prickly sculpin (*Cottus asper*) are a benthic intraguild predator, and their presence selects for a more limnetic stickleback ecotype [46]. We therefore collected data on the presence/absence of prickly sculpin in all sampling locations from Hutchinson et al. [64], Miller et al. [48], Atkinson [65], Dennenmoser et al. [66] and Vamosi [61] (see Supplementary Material for data sources for each lake).

The pH [67] and dissolved calcium concentrations [68] of lake water have previously been associated with external bony armour in stickleback (a trait which varies between benthic and limnetic ecotypes). Therefore we also measured these variables, the former with a calibrated pH meter (Multi 340i, WTW, Weilheim, Germany) and the latter were obtained by collecting two filtered water samples (one acidified with 2% nitric acid, one frozen) from each lake. The dissolved calcium concentration (to the nearest mg/L) was then measured from the water samples at the Division of Agriculture & Environmental Science at the University of Nottingham by inductively coupled plasma mass spectrometry (ICP-MS).

### Stickleback sampling

Stickleback were caught using unbaited minnow traps set overnight from the lake shores during spring of 2015 (all stickleback ecotypes move to the shallows during the spring to breed). Samples of between 10 and 63 individuals (See Table S1 for lake specific sample sizes) were taken from each lake and transported to a rental property in aerated lake water for processing. Immediately prior to processing, fish were euthanised with an overdose of tricaine methanesulfonate (‘MS222’) (400 mg L^-1^), and killed by destruction of the brain, in accordance with Schedule One of UK Home Office regulations and with the approval of the University of British Columbia Animal Care Committee (UBC animal care certificate A11-0402). Fin clips were immediately taken and stored in 90% Ethanol for later genomic analyses.

### Identification of benthic-limnetic divergence

#### Phenotypic quantification

Fish sampled from lakes containing species pairs (PAXT and PRIE) were visually classified as benthic or limnetic at the time of capture as well as being later measured for all phenotypic traits.

To determine body size, fish were blotted and weighed to the nearest milligram. To assess body shape differences, each stickleback’s left side was photographed using a tripod mounted digital SLR camera fitted with a macro lens and macro digital ring flash. Images were scaled, and 13 landmarks were placed on each image using tpsDig, version 2.16 [69]. Landmark data were then exported to MorphoJ, version 1.06d [70]. A Procrustes fit was performed to align specimens by their main axes and remove size and rotation bias. Differences between lakes were identified using a Procrustes ANOVA with lake as the classifier. Allometric variation in body shape was removed by taking the residuals of a multivariate partial least squares regression against log centroid size, and the regression was pooled within lakes because the Procrustes ANOVA indicated differences between group centroids [71]. Regression residuals were exported into R, version 3.5.2 [72], where they were standardised and scaled, and variation in body shape was reduced to a single axis using a principal components analysis (PCA), implemented by singular value decomposition. This principal axis (shape PC1) was used to describe differences in body shape in all further analyses.

To assess differences in body armour, fish were first bleached and then stained with alizarin red to highlight external skeletal structures following standard procedure [73]. Fish were then re-photographed as above, images were scaled, and counts of lateral plate number, alongside measurements of standard length, first and second dorsal spine length, longest plate length, pelvis height, pelvis length and pelvic spine length, were taken (continuous elements to the nearest 0.01mm) using ImageJ, version 1.52a [74]. All continuous armour variables (thus excluding plate number, which was independent of body size in our data set) were size-standardized by taking the residuals of a regression against standard length. Body armour variables were highly correlated, thus we used a principal components analysis (PCA) to reduce variation in body armour variables to a single axis: armour PC1. Armour PC1 was used to describe differences in body armour in all further analyses.

Finally, the left primary gill arch was extracted from each individual. For each gill arch, the total number of gill rakers were counted, and the mean gill raker length was calculated by taking the mean of the length of the first three rakers on each arch, measured to the nearest micrometre.

### Genomic SNP analyses

DNA was isolated from fin tissue using Quiagen Blood and Tissue DNA purification kits. RAD-seq data was generated following Magalhaes et al. [75]. BAM files were produced following Magalhaes et al. [75]. Variants were called from per-individual BAM files to create a single VCF file using the Stacks pipeline [76] in Stacks, version 1.47. The POPULATIONS program in Stacks was run with the following filters: SNPs with a minimum depth of coverage < 3 were removed; SNPs present in < 50% of individuals within a population were removed; SNPs with a minor allele frequency < 0.05 were removed; and SNPs that were not present in all of the populations were removed. VCFtools, version 0.1.16 [77], was then used to remove sites with mean depth values (over all individuals) < 6 and > 200, sites with > 25% missing data, sites with a minor allele count over all individuals < 2 and the sex chromosomes. This pipeline produced an overall dataset of 12,756 SNPs for 333 individuals across the 21 lakes (dataset 1). This dataset was then subject to further filtering for some analyses, and detailed information about individual RAD datasets is given in Table 1.

### Linkage disequilibrium

Sets of loci that have a tendancy to be inherited together, and thus are highly correlated, tend to be affected by the same evolutionary processes and so contain useful information for identifying the characteristics of the processes affecting each set of linked loci e.g. whether divergence is likely linked to small genomic regions e.g. inversions, or is genome wide. To investigate whether any groups of linked loci would distinguish the genomic clusters identified in other genomic analyses we performed a linkage disequilibrium network analysis (LDna) using the LDna package in R. The r^2^ linkage disequilibrium matrix was generated using dataset 1 (12,756 SNPs) in Plink version 1.9 [78]. For the extractClusters step of LDna the minimum number of edges was set to 100 and Φ was set to five. SNPs in each LD cluster were extracted from dataset 1 using VCFtools, VCF files were read into R using the vcfR package and principal components anaylsis (PCA) of the SNPs in each LD cluster was performed using the adegenet [79] package.

Many genomic tools, however, rely on the assumption that variants are independent and therefore SNPs in linakge diequilibrium must be removed for such analyses. To that end, we estimated linkage disequilibrium across the genome as a whole by calculating pairwise R^2^ values in 100kb sliding windows using Plink version 1.9. R^2^ values range between 0 (no linkage) and 1 (complete linkage), and therefore a relatively conservative LD threshold was set at R^2^ > 0.2. Thinning dataset 1 (12,756 SNPs) to unlinked loci resulted in a dataset with 9,668 retained SNPs (dataset 2, Table 1).

### Genomic patterns

We used fineRADstructure [33] to construct a co-ancestry matrix using the primary SNP set including all 333 individuals (dataset 1, 12,756 SNPs). Prior filtering for linkage disequilibrium is not necessary for analyses using the RADpainter tool at it efficiently estimates the effective number of loci in mapped data files during the analysis [33]. The fineSTRUCTURE [80] clustering algorithm was run with a burn-in of 100,000 iterations followed by 100,000 sampled iterations and the tree building algorithm was run with a burn-in of 10,000 iterations. We then performed a principal coordinate analysis (PCoA) on the linkage filtered dataset (dataset 2, 9668 SNPs). The PCoA was performed using Elucidean distances with the package adegenet in R. VCF files were converted to genpop format for input to adegenet using PGDSpider, version 2.1.1.5 [81].

### Phenotypic divergence

Genomic analyses grouped all fish into two broad genomic clusters, cluster 1 and cluster 2. Although marine fish were grouped with the freshwater fish in cluster 1, they were treated as a third, separate, group in all further analyses. Additionally, they were excluded from most subsequent genetic analyses because their presence in freshwater/coastal areas is transient (they migrate to shallow coastal areas only in the spring to breed) and they represent the likely ancestral state of all freshwater populations [82]. To determine the phenotype of these three groups (marine, cluster 1 and cluster 2), we calculated the mean of each phenotypic variable (weight, number of gill rakers, mean raker length, armour PC1 and shape PC1) for each group. To test whether the means of each phenotypic variable in each of the three groups were significantly different from one another, linear mixed models were performed using the nlme package [83], with lake included as a random effect and group (marine, cluster 1, cluster 2) as a fixed effect. For models showing a significant effect of group, post-hoc pairwise comparisons were performed using estimated marginal means, implemented using the emmeans package [84] in R. P-values for post-hoc comparisons were adjusted for multiple testing using the False Discovery Rate (FDR) method [85].

### Phylogenetic analyses

Prior to phylogenetic analysis, we filtered our master dataset (dataset 1) to remove all known QTL in stickleback. Data for QTL were downloaded from Peichel and Marques [59], converted to BED format, and removed from the VCF file using VCFtools. This reduced the number of SNPs from 12,756 to 8351. We then ensured approximate linkage equilibrium of remaining markers by removing all SNPs with an R^2^ value >0.2 using Plink version 1.9. This left 6215 SNPs, dataset 3.

To construct a phylogeny of all sequenced individuals, we used a bootstrapped maximum-likelihood based approach, implemented in RAxML, version 8.2.12 [86]. The VCF file was converted to phylip format for input to RAxML using python version 3.8.2. RAxML was run with a GTR-GAMMA model of substitution rate heterogeneity, automatic bootstrap replicate halting using the autoMRE function and with the default settings for all other parameters.

To assess the robustness of the maximum likelihood phylogeny, we also performed topology weighting using TWISST [87]. Topology weighting was carried out on four freshwater populations, with a single marine sequence (from LICA) as the outgroup. The freshwater populations were selected to contain two pairs of geographically proximal populations, with one pair from either side of the Georgia strait (NORT and BEAV, and DOUG and KLEN), and with one population from each pair falling in cluster 1, the other in cluster 2. Dataset 2 (9668 SNPs) was filtered to contain all individuals from each of the four freshwater populations and a single individual from LICA (TWISST only accepts a single sequence as an outgroup), using VCFtools (dataset 4, 9668 SNPs). The VCF file was converted to .geno format and maximum likelihood trees were estimated in phyml [88] in sliding windows of 50bp using Python 2.7.15 and the scripts available with TWISST. Topology weightings were then computed using Python 3.8.2 and topologies were visualised in R.

### Phylogenetic signal in phenotypic traits

To estimate phylogenetic signal, the phylogeny constructed in RAxML was imported into R using the ape package [89], individual nodes were collapsed to leave a single node per population, with the exception of the two marine populations, which were both collapsed into one node using the phytools [90] and phangorn [91] R packages. Phenotypic trait data (weight, number of gill rakers, mean raker length, armour PC1 and shape PC1) were added to the tree tips, and phylogenetic signal and associated *p*-values for each trait were estimated using the package phylosignal [92]. We used Pagel’s λ [93] to estimate phylogenetic signal as this statistic performs well compared to others available and has a low type 1 error rate [39, 94, 95]. *P*-values are calculated using likelihood ratio tests that compare the observed λ statistic with a phylogenetically independent trait distribution. We also simulated data for five additional traits to be distributed randomly with regard to phylogeny. Simulated traits were tested alongside the real phenotypic variables for comparison.

As we aim to detect whether benthic and limnetic characteristics have evolved a single time or repeatedly across the radiation, we also used the R package SURFACE [32] to estimate the most likely number of different selection regimes (*k*) and instances of convergent evolution (*c*) by identify the best fitting model of trait evolution for our phylogeny and associated phenotypic traits. SURFACE begins by fitting a single peak Ornstein-Uhlenbeck (OU) model (which allows for a single adaptive optimum and variation in the parameter α, which describes the strength of selection towards that optimum) by maximum likelihood. It then sequentially adds adaptive peaks to the model in a step-wise process and accepts each more complex model until AIC values are no longer improved. SURFACE then attempts to collapse regimes with the same optima in a process of step-wise backwards selection whereby if multiple optima are the same, the AIC of the model is improved by reducing the number of model parameters. We ran SURFACE using the same collapsed phylogeny and associated trait data that was generated to estimate phylogenetic signal. The tree was converted to ouchtree format and the best fitting model of trait evolution was estimated under an AIC threshold of 0 (any improvement in AIC should be accepted) using the SURFACE R package.

### Relationship between phenotype and environment

To investigate associations between environmental characteristics and divergence in phenotypic traits, we used a phylogenetic generalised least squares (PGLS) approach so that phylogenetic signal could be accounted for in the models, using the ape [89], nlme [96] and geiger [97] packages in R. Marine fish were excluded from all phenotype – environment modelling because our main aim was to detect effects in relation to the freshwater benthic and limnetic phenotypes in BC and, although marine fish have a limnetic phenotype, we found them to differ phenotypically from the freshwater limnetic fish in BC. Separate models were run for each phenotypic trait (weight, number of gill rakers, mean gill raker length, armour PC1 and shape PC1). Models were fitted by maximum likelihood and we began with all environmental variables in each model (mean lake depth [m], lake area [km^2^], presence/absence of prickly sculpin, pH and calcium concentration [mg/L]). Terms were then removed sequentially, with the least significant terms removed first, until the reduced model was no longer a significant improvement on the fuller model under the *p*<0.05 threshold. Model comparison was conducted using Wald tests. Phylogenetic effects for each phenotypic trait were accounted for in each model following the principles set out in Mazel et al. [98]. We first transformed the phylogeny for each phenotypic trait under a lambda model with lambda specified as the lambda estimate for that phenotypic trait in the phylogenetic signal analyses. The phylogenetic variance-co-variance matrices were computed from the transformed trees using the ape package and converted to correlation matrices, which were used to specify phylogenetic correlation of errors in the models.

## Supplementary Tables

**Table S1.**
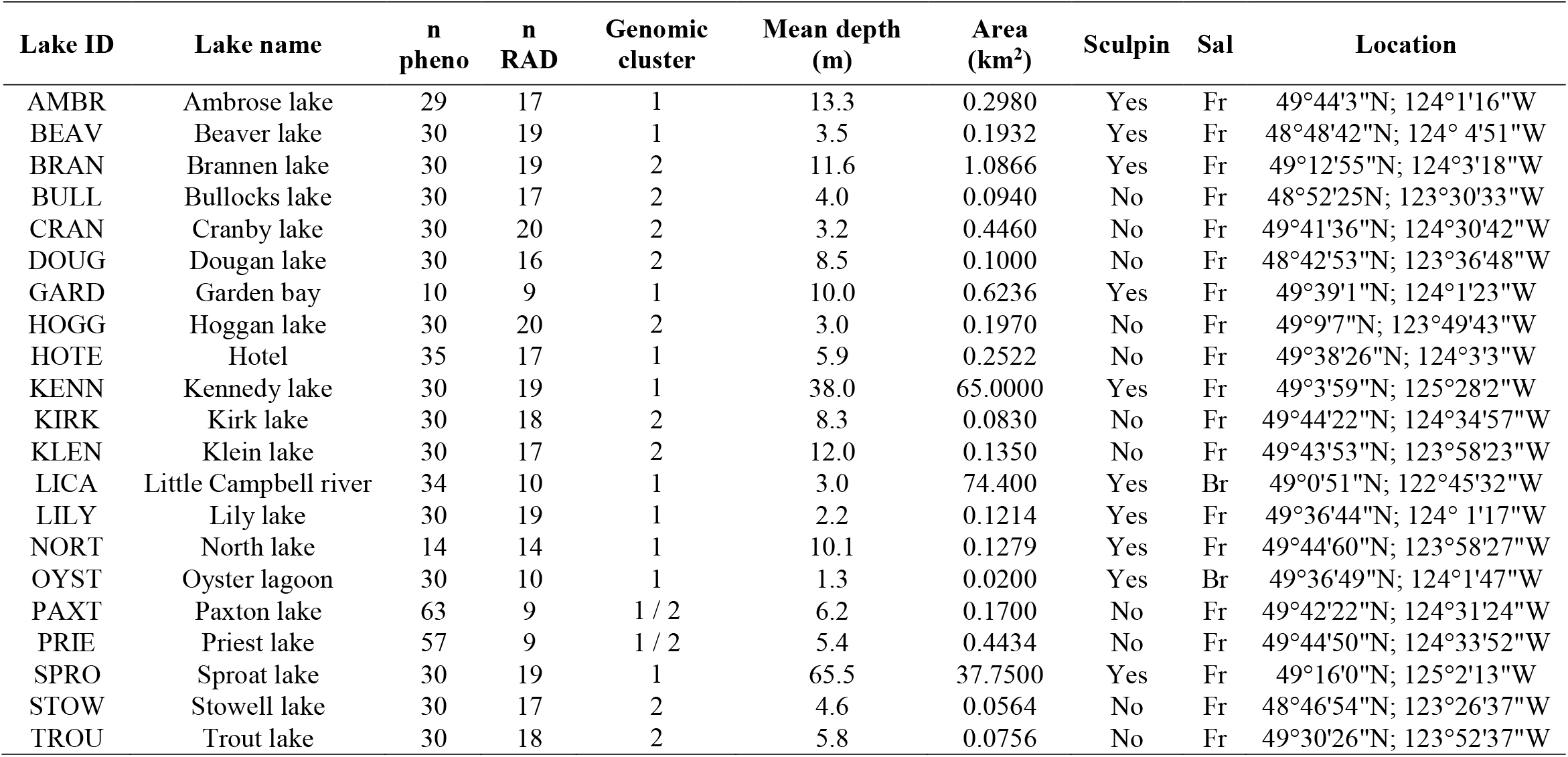
Description of sample sites. Sample sizes are shown for phenotypic (n pheno) and RAD (n RAD) analyses. Sal refers to salinity classifications, Fr: freshwater (absolute conductivity <500 μS/cm), Br: brackish (absolute conductivity 20,000-35,000 μS/cm). Sampling locations are given by latitude followed by longitude.

| Lake ID | Lake name | n<br>pheno | n<br>RAD | Genomic<br>cluster | Mean depth<br>(m) | Area<br>(km<sup>2</sup>) | Sculpin | Sal | Location |
| --- | --- | --- | --- | --- | --- | --- | --- | --- | --- |
| AMBR | Ambrose lake | 29 | 17 | 1 | 13.3 | 0.2980 | Yes | Fr | 49°44'3"N; 124°1'16"W |
| BEAV | Beaver lake | 30 | 19 | 1 | 3.5 | 0.1932 | Yes | Fr | 48°48'42"N; 124° 4'51"W |
| BRAN | Brannen lake | 30 | 19 | 2 | 11.6 | 1.0866 | Yes | Fr | 49°12'55"N; 124°3'18"W |
| BULL | Bullocks lake | 30 | 17 | 2 | 4.0 | 0.0940 | No | Fr | 48°52'25"N; 123°30'33"W |
| CRAN | Cranby lake | 30 | 20 | 2 | 3.2 | 0.4460 | No | Fr | 49°41'36"N; 124°30'42"W |
| DOUG | Dougan lake | 30 | 16 | 2 | 8.5 | 0.1000 | No | Fr | 48°42'53"N; 123°36'48"W |
| GARD | Garden bay | 10 | 9 | 1 | 10.0 | 0.6236 | Yes | Fr | 49°39'1"N; 124°1'23"W |
| HOGG | Hoggan lake | 30 | 20 | 2 | 3.0 | 0.1970 | No | Fr | 49°9'7"N; 123°49'43"W |
| HOTE | Hotel | 35 | 17 | 1 | 5.9 | 0.2522 | No | Fr | 49°38'26"N; 124°3'3"W |
| KENN | Kennedy lake | 30 | 19 | 1 | 38.0 | 65.0000 | Yes | Fr | 49°3'59"N; 125°28'2"W |
| KIRK | Kirk lake | 30 | 18 | 2 | 8.3 | 0.0830 | No | Fr | 49°44'22"N; 124°34'57"W |
| KLEN | Klein lake | 30 | 17 | 2 | 12.0 | 0.1350 | No | Fr | 49°43'53"N; 123°58'23"W |
| LICA | Little Campbell river | 34 | 10 | 1 | 3.0 | 74.400 | Yes | Br | 49°0'51"N; 122°45'32"W |
| LILY | Lily lake | 30 | 19 | 1 | 2.2 | 0.1214 | Yes | Fr | 49°36'44"N; 124° 1'17"W |
| NORT | North lake | 14 | 14 | 1 | 10.1 | 0.1279 | Yes | Fr | 49°44'60"N; 123°58'27"W |
| OYST | Oyster lagoon | 30 | 10 | 1 | 1.3 | 0.0200 | Yes | Br | 49°36'49"N; 124°1'47"W |
| PAXT | Paxton lake | 63 | 9 | 1 / 2 | 6.2 | 0.1700 | No | Fr | 49°42'22"N; 124°31'24"W |
| PRIE | Priest lake | 57 | 9 | 1 / 2 | 5.4 | 0.4434 | No | Fr | 49°44'50"N; 124°33'52"W |
| SPRO | Sproat lake | 30 | 19 | 1 | 65.5 | 37.7500 | Yes | Fr | 49°16'0"N; 125°2'13"W |
| STOW | Stowell lake | 30 | 17 | 2 | 4.6 | 0.0564 | No | Fr | 48°46'54"N; 123°26'37"W |
| TROU | Trout lake | 30 | 18 | 2 | 5.8 | 0.0756 | No | Fr | 49°30'26"N; 123°52'37"W |

**Table S2.**
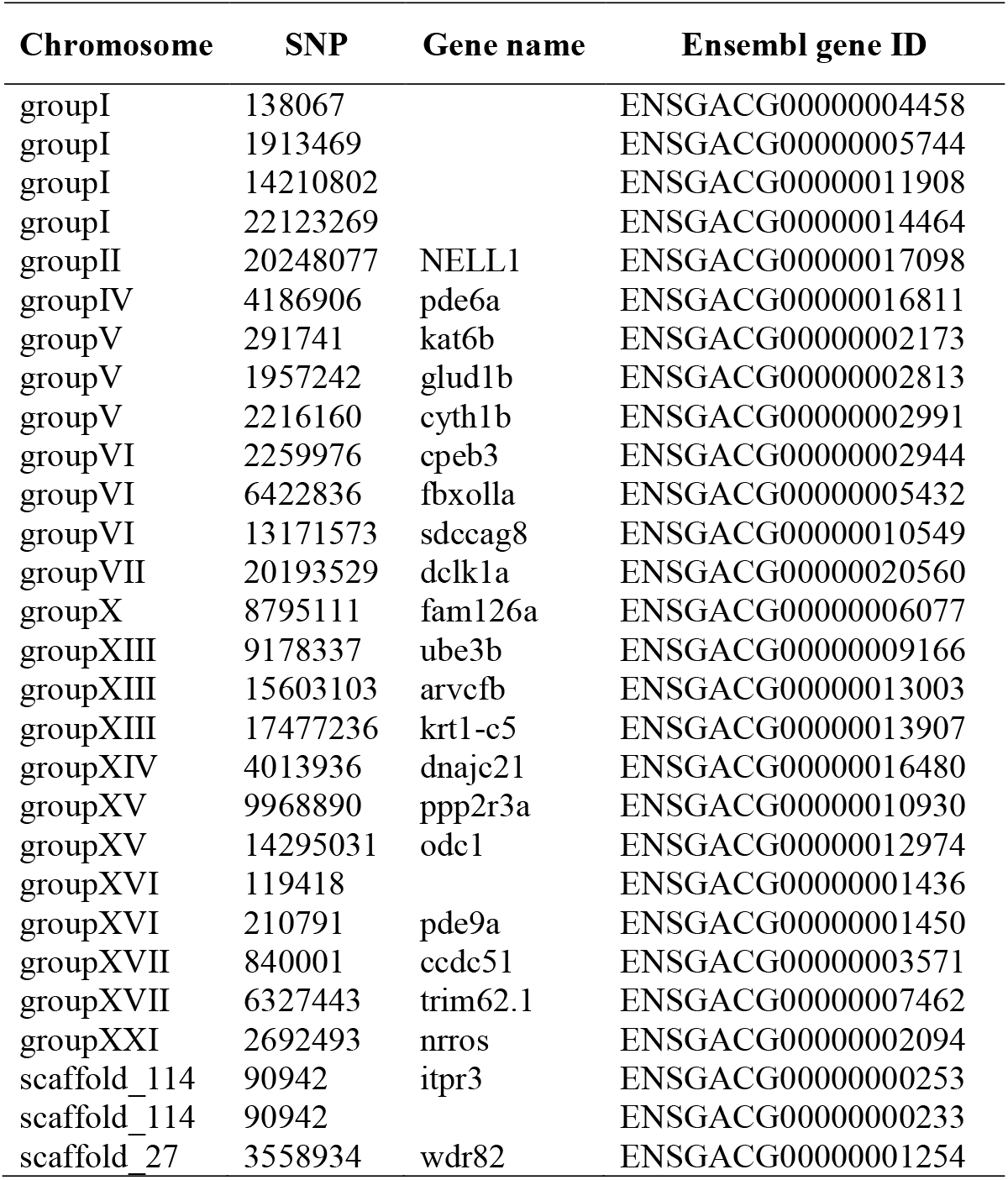
SNPs in linkage cluster 10 that lie within genes. Genome location of SNPs falling within coding regions and the associated gene and gene ID for each SNP.

**Table S3.**
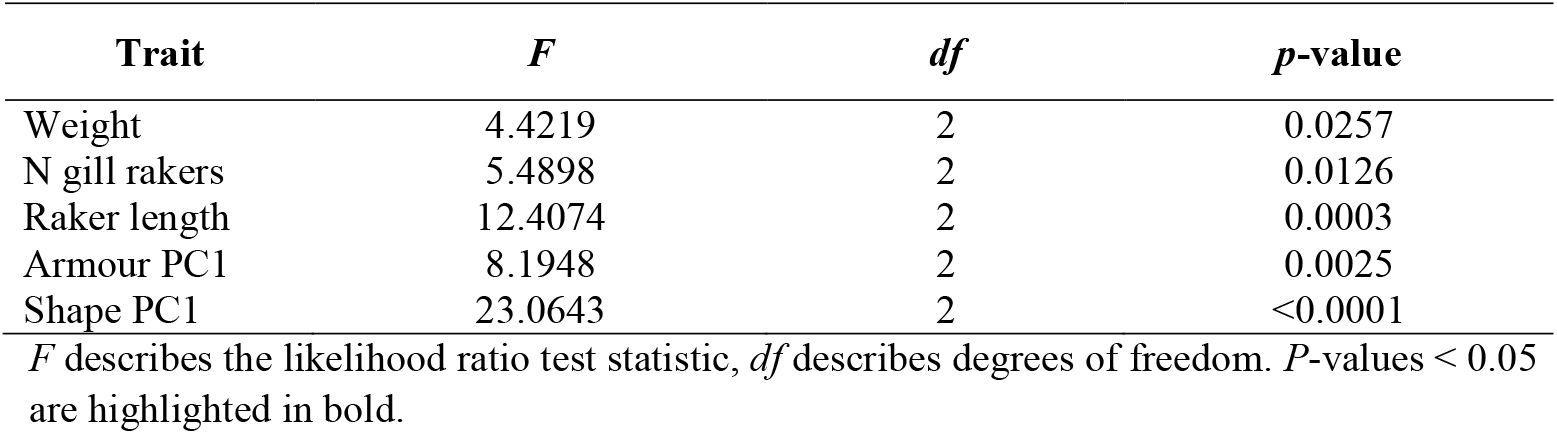
Linear mixed model results for phenotypic differences between groups. Table shows the results of linear mixed models testing for differences between three groups (marine, genomiccluster 1 and genomic cluster 2) in five phenotypic traits, with population as a random effect.

**Table S4.** Post-hoc Estimated marginal means test results for pairwise phenotypic differences between groups. Table shows the results of post-hoc estimated marginal means tests to determine pairwise differences between groups (marine, genomic cluster 1 and genomic cluster 2) when phenotypes differed significantly between groups in linear mixed models.

| Trait | Groups compared | Estimate | SE | df | Adjusted *p*-value |
| --- | --- | --- | --- | --- | --- |
| Weight | limnetic-like – benthic-like | 0.2453 | 0.2400 | 20 | 0.3188 |
|  | marine – benthic-like | -1.0119 | 0.4193 | 20 | **0.0383** |
|  | marine – limnetic-like | -1.2572 | 0.4232 | 20 | **0.0227** |
| N gill rakers | limnetic-like – benthic-like | -2.0254 | 0.6387 | 20 | **0.0144** |
|  | marine – benthic-like | -1.9978 | 1.1211 | 20 | 0.1349 |
|  | marine – limnetic-like | 0.0276 | 1.1304 | 20 | 0.9808 |
| Raker length | limnetic-like – benthic-like | -0.1799 | 0.0639 | 20 | **0.0107** |
|  | marine – benthic-like | -0.5286 | 0.1121 | 20 | **0.0004** |
|  | marine – limnetic-like | -0.3487 | 0.1130 | 20 | **0.0088** |
| Armour PC1 | limnetic-like – benthic-like | -2.2551 | 0.6914 | 20 | **0.0069** |
|  | marine – benthic-like | -3.8791 | 1.2155 | 20 | **0.0069** |
|  | marine – limnetic-like | -1.6240 | 1.2250 | 20 | 0.1999 |
| Shape PC1 | limnetic-like – benthic-like | -2.4654 | 0.4697 | 20 | **<0.0001** |
|  | marine – benthic-like | -4.5652 | 0.8202 | 20 | **<0.0001** |
|  | marine – limnetic-like | -2.0998 | 0.8280 | 20 | **0.0197** |

Estimates describe the mean difference between groups, SE describes the standard error of the estimates and *df* describe degrees of freedom. *P*-values are adjusted for multiple testing using the FDR method. *P*-values < 0.05 are highlighted in bold.

**Table S5.** Effects of the environment on the distribution of benthic-limnetic phenotypic traits. Table shows the results of phylogenetic generalised least squares analyses (PGLS) on each phenotypic trait. *P*-values < 0.05 are highlighted in bold. *P*-values were adjusted for multiple testing using the FDR method.

| Phenotypic trait | Environmental variable | df | Wald statistic | *p*-value | Adjusted *p*-value |
| --- | --- | --- | --- | --- | --- |
| Armour PC1 | Lake area | 1 | 1.5877 | 0.2077 | 0.5077 |
|  | Ca concentration | 1 | 3.2274 | 0.0724 | 0.2172 |
|  | Mean lake depth | 1 | 0.0883 | 0.7663 | 0.8969 |
|  | pH | 1 | 1.6609 | 0.1975 | 0.3950 |
|  | Sculpin presence | 1 | 17.6411 | **0.0000** | **0.0002** |
| N rakers | Lake area | 1 | 0.0461 | 0.8300 | 0.8300 |
|  | Ca concentration | 1 | 0.3525 | 0.5527 | 0.6088 |
|  | Mean lake depth | 1 | 0.0327 | 0.8564 | 0.8969 |
|  | pH | 1 | 0.1860 | 0.6663 | 0.7882 |
|  | Sculpin presence | 1 | 0.7284 | 0.3934 | 0.4721 |
| Raker length | Lake area | 1 | 6.8857 | **0.0087** | 0.0521 |
|  | Ca concentration | 1 | 1.4304 | 0.2317 | 0.4634 |
|  | Mean lake depth | 1 | 0.0168 | 0.8969 | 0.8969 |
|  | pH | 1 | 0.5091 | 0.4755 | 0.7133 |
|  | Sculpin presence | 1 | 2.6932 | 0.1008 | 0.1512 |
| Shape PC1 | Lake area | 1 | 0.0641 | 0.8001 | 0.8300 |
|  | Ca concentration | 1 | 0.2620 | 0.6088 | 0.6088 |
|  | Mean lake depth | 1 | 0.1346 | 0.7137 | 0.8969 |
|  | pH | 1 | 1.7163 | 0.1902 | 0.3950 |
|  | Sculpin presence | 1 | 3.8490 | **0.0498** | 0.0996 |
| Weight | Lake area | 1 | 0.1549 | 0.6939 | 0.8300 |
|  | Ca concentration | 1 | 0.5770 | 0.4475 | 0.6088 |
|  | Mean lake depth | 1 | 16.8205 | **0.0002** | **0.0013** |
|  | pH | 1 | 7.9713 | **0.0048** | **0.0285** |
|  | Sculpin presence | 1 | 0.0013 | 0.9714 | 0.9714 |

## Supplementary Figures

**Figure S1.**
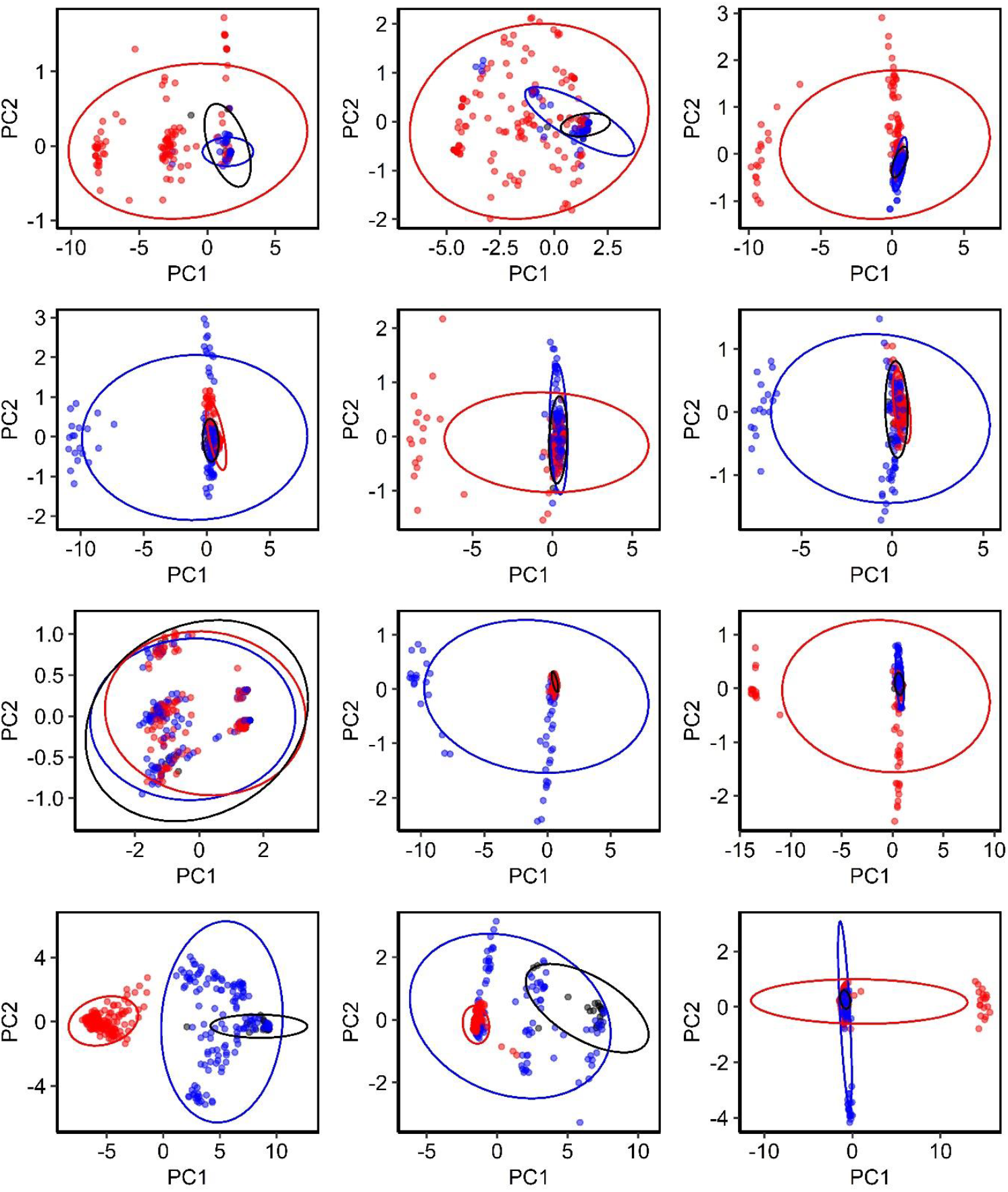
Principal component analyses (PCA) of each LD cluster in the LD network identified by LDna. Black circles represent marine individuals, blue circles, freshwater cluster 1 and red circles freshwater cluster 2. 95% confidence ellipses are shown for each PCA in the corresponding colours.

**Figure S2.**
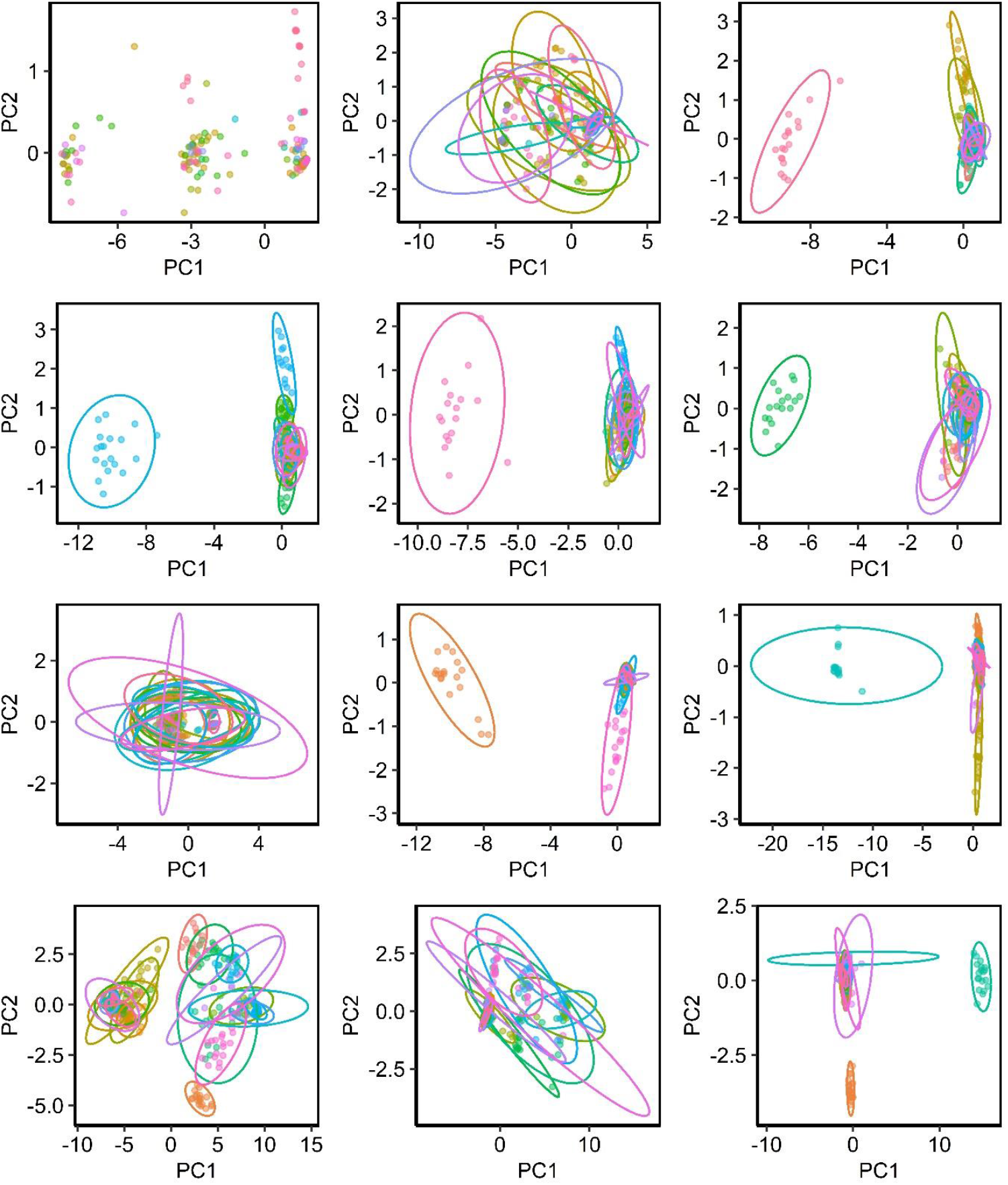
Principal component analyses (PCA) of each LD cluster in the LD network identified by LDna. Coloured circles and ellipses represent each of the 23 sampled populations.

